# HTRF-based identification of small molecules targeting SARS-CoV-2 E protein interaction with ZO-1 PDZ2

**DOI:** 10.1101/2025.07.16.665136

**Authors:** Flavio Alvarez, Florence Larrous, Ariel Mechaly, Benjamin Bardiaux, Quentin Goor, Ahmed Haouz, Blandine Seon-Meniel, Rodolphe ALVES De Sousa, Stéphane Bourg, Bruno Figadere, Nicolas Wolff, Hélène Munier-Lehmann, Célia Caillet-Saguy

## Abstract

The SARS-CoV-2 E protein through its C-terminal PDZ-binding motif (PBM) interacts with several host PDZ-containing proteins, including the ZO-1 protein via its PDZ2 domain, thereby contributing to viral pathogenesis. Targeting this interaction represents a potential therapeutic strategy.

In this study, we determined the X-ray structure of the E PBM peptide in complex with the ZO-1 PDZ2 domain at 1.7 Å resolution. The structure revealed a domain-swapped dimer conformation of ZO-1 PDZ2, with the E PBM peptide conventionally bound within the PDZ domain’s canonical binding groove exhibiting key interactions characteristic of type II PBM–PDZ interactions.

To identify potential inhibitors of the E PBM/ZO-1 PDZ2 interaction, we performed a HTRF screening using a protein-protein interaction-focused library of 1,000 compounds. This led to the identification of 36 hits that disrupted this interaction. Subsequent cytotoxicity and dose-response assays narrowed the selection to 14 promising compounds.

Docking simulations showed that some compounds bind within or near the PBM-binding pocket, supporting a competitive mechanism of interaction inhibition, while others bind at a central interface between the two PDZ monomers, suggesting an inhibition of dimerization, which in turn prevents PBM binding. Thus, the ZO-1 PDZ2–E PBM interaction can be inhibited through both direct and indirect mechanisms.

Finally, antiviral assays using a NanoLuciferase-expressing recombinant SARS-CoV-2 demonstrated that one compound, C19, significantly reduced viral replication, highlighting its potential as a candidate for further therapeutic development.

**Highlights:**

- Crystal structure reveals the binding determinant of E protein PBM to ZO-1 PDZ2 dimer
- HTRF screening identified 36 inhibitors of the E–ZO-1 protein interaction
- Docking revealed dual mechanisms: PBM groove binding and dimer interface disruption
- Compound C19 reduced SARS-CoV-2 replication in a NanoLuc reporter assay
- Targeting E–ZO-1 interaction may prevent barrier loss and systemic COVID-19 effects

**Graphical abstract:** 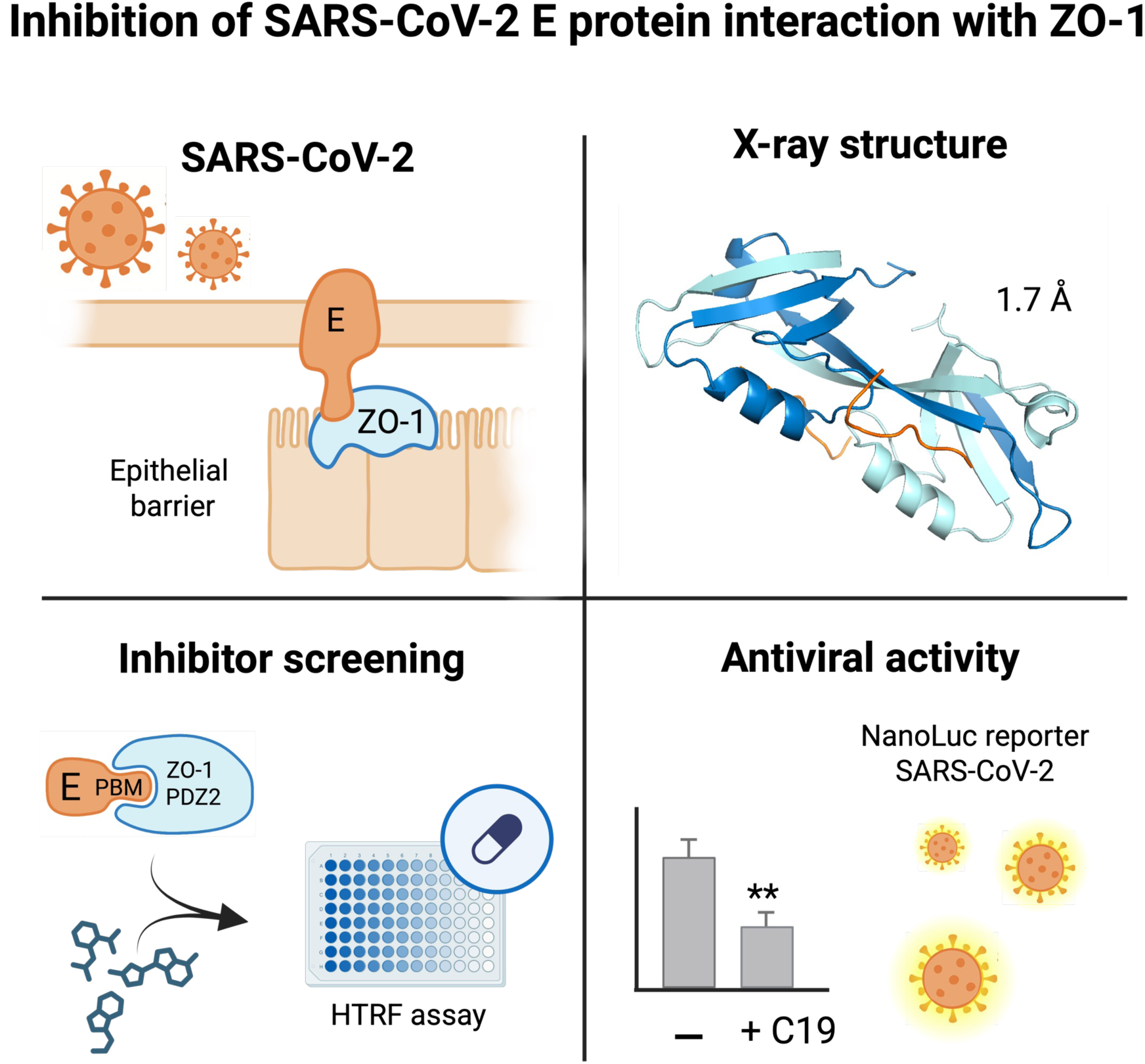

## Introduction

The SARS-CoV-2 virus continues to pose a significant public health challenge due to its ongoing mutations and the emergence of new variants, which contribute to periodic outbreaks worldwide ^1,2^. SARS-CoV-2 can lead to severe respiratory illness in humans, commonly referred to as COVID-19, often characterized by a cytokine storm leading to acute respiratory distress syndrome (ARDS) and high mortality rates ^3–5^. Current FDA-approved treatments primarily target viral proteins essential for replication, including antiviral drugs such as Paxlovid, Remdesivir, and Molnupiravir ^6^. These treatments remain effective, targeting highly conserved viral regions such as the SARS-CoV-2 main protease (Mpro) and the viral RNA-dependent RNA polymerase. However, monoclonal antibodies targeting the spike protein, another widely used therapeutic strategy, have shown limitations ^7–9^. Their effectiveness decreases as spike protein mutations emerge, highlighting the urgent need for alternative therapeutic targets.

SARS-CoV-2 is an enveloped, positive-sense single-stranded RNA virus that encodes several structural and non-structural proteins crucial for its viral life cycle. Among the structural proteins, the envelope (E) protein stands out for its pivotal role in viral assembly, release, and host-pathogen interactions ^10,11^. This small and highly conserved integral membrane protein acts as a viroporin, enabling ion transport across host cell membranes, and contributes to viral morphogenesis. Notably, the E protein has an exceptionally low mutation rate, with a conservation rate exceeding 99.98%, suggesting its critical role in viral fitness ^12,13^.

Given its conserved nature and functional importance in viral assembly and pathogenesis, the SARS-CoV-2 E protein has emerged as an attractive therapeutic target. Recent studies, mainly through computational approaches, have identified natural and synthetic compounds capable of inhibiting its viroporin activity or interactions with host proteins. For instance, compounds derived from flavonoids, such as rutin, and glycyrrhizic acid, a triterpene glycoside, have demonstrated the ability to bind and destabilize the E protein ^12^.

Therapeutic approaches targeting protein-protein interactions (PPIs) involving the E protein remain largely underexplored, especially as its interactome discoveries continue to expand ^14–18^. The E protein features a short C-terminal interaction motif, the PDZ binding motif (PBM), capable of interacting with host cellular PDZ domain-containing proteins, leading to the disruption of endogenous PPIs involved in various cellular functions. The PBM of the SARS-CoV-2 E protein and its interactions with host proteins have been studied ^15,16,19–21^. We and others have shown that the PBM plays a significant role in viral pathogenicity ^22,23^. Our previous work demonstrated that the PBM is essential for SARS-CoV-2 virulence, with its underlying molecular mechanism intricately linked to its interaction with ZO-1, a key factor in COVID-19 pathogenesis ^22,24^. Therefore, the E protein interacts with multiple host cell proteins critical for cell polarity and epithelial integrity, including the tight junction proteins PALS-1 and ZO-1 ^15,20,24–26^. These interactions weaken tight junctions and epithelial barriers increasing the permeability of lung tissue and other organs, eventually leading to fluid leakage ^27,28^. This contributes to severe complications such as pulmonary edema and ARDS, which are commonly observed in critical COVID-19 cases. Additionally, the breakdown of tight junctions facilitates the uncontrolled migration of immune cells and inflammatory mediators into the affected tissues, amplifying the immune response and potentially triggering a cytokine storm, a major factor in the severity of COVID-19 ^29,30^. In this context, targeting the interaction between the E protein and ZO-1 may represent a novel therapeutic approach to limit viral propagation and mitigate the pathogenesis and severity of the disease.

In this study, we aimed to identify small molecules capable of inhibiting the interaction between the SARS-CoV-2 E protein and the PDZ2 domain of the ZO-1 protein. Using a Homogeneous Time-Resolved Fluorescence (HTRF) assay, we screened a PPI-focused library of 1,000 small compounds ^31^ and identified 36 candidates for further validation. Inhibition and cytotoxicity dose-response assays allowed us to select 14 promising candidates. Their antiviral activities were assessed in vitro using a recombinant SARS-CoV-2 containing a NanoLuciferase reporter gene. One of these molecules demonstrated significant antiviral activity. We solved the X-ray structure of the complex between the PDZ2 of ZO-1 and the PBM of the E protein and performed docking studies with selected inhibitors to elucidate their mode of action. Our findings provide insights into novel therapeutic strategies targeting the interaction between the SARS-CoV-2 E protein and the PDZ-containing ZO-1 protein. This approach could help maintain barrier integrity, alleviate inflammation, reduce the severity of COVID-19, and potentially provide a groundbreaking therapeutic strategy to combat other viruses employing similar mechanisms.

## Results

### The crystallographic structure reveals key interaction determinants between the ZO-1 PDZ2 swapped dimer and the C-terminal region of the E protein

To gain structural insights into the interaction between ZO-1 and the C-terminal PBM of the SARS-CoV-2 E protein, we solved the X-ray structure of the ZO-1 PDZ2 domain in complex with the last 12-residue peptide of the E protein (_ac_NLNSSRVPDLLV_COOH_). The crystal structure of ZO1- PDZ2 in complex with the SARS-CoV-2-E-PBM peptide was resolved at 1.72 Å using the molecular replacement method (**Figure 1**, **Table 1**). The final refined model contains two PDZ domains per asymmetric unit, adopting a swapped dimer conformation. Each PDZ domain features a fold comprising five β strands and two α helices. Notably, the N-terminal region, encompassing strands β1 and β2, does not contribute to the core PDZ fold of its own molecule. Instead, these segments complete the PDZ fold of the adjacent molecule in a reciprocal manner. Two primary regions of intermolecular interactions between the PDZ domains significantly contribute to the extensive dimer interface: the inter-domain β1/β6 pairing and the extended antiparallel β2- β3/β2’-β3’ assembly (**Figure 1A**).

**Figure 1.**
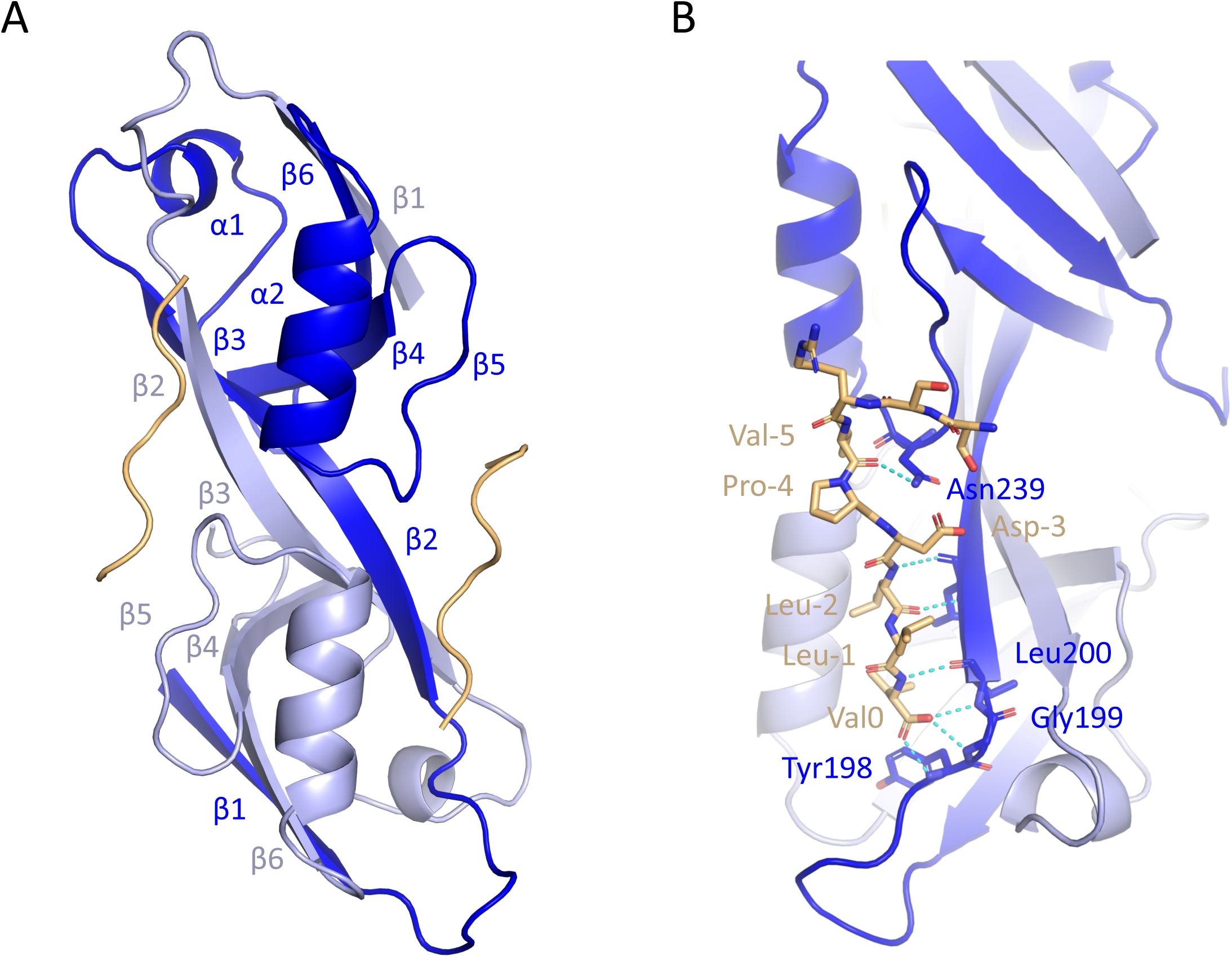
X-ray crystal structure of the PDZ2 domain of ZO-1 bound to the SARS-CoV-2 E protein PBM. A. Ribbon diagram of the final crystal structure model of the ZO-1 PDZ2 domain in complex with the SARS-CoV-2 E protein PBM peptide. The structure forms a swapped dimer, with one monomer colored light blue and the other dark blue. The bound peptides are shown in orange. Secondary structure elements are labeled according to the canonical PDZ domain scheme. B. Close-up view of the SARS-CoV-2 E protein PBM bound within the PDZ domain. Key interacting residues are shown as sticks and labeled. Intermolecular hydrogen bonds and polar interactions are indicated by cyan dashed lines.

**Table 1.** Data collection and refinement statistics. Statistics for the highest-resolution shell are shown in parentheses.

The swapped dimer assembly observed in the ZO-1 PDZ2 complex was similarly identified for ZO-1 and ZO-2 PDZ2 ^32^ either in their ligand-free forms or in complex with the C-terminal peptide of connexin-43 ^33^ (**Supplementary Figure 1**). Two peptides are bound to each PDZ dimer (**Figure 1A**). Electron density maps suggest a well-defined conformation of the last eight residues of the peptide containing the PBM. The E PBM binds to the PDZ in a conventional manner, inserting into a binding groove formed by the β2 strand, the α2 helix, and the “GLGF” motif, and extending the antiparallel β-sheet. The root-mean-square deviation (RMSD) between the structures of ZO-1 PDZ2 in complex with connexin-43 and the E protein peptides is 0.54 Å, indicating a high degree of structural similarity (**Supplementary Figure 1**). The last six residues of the SARS-CoV-2-E-PBM peptide are in contact with the PDZ domain, whereas the upstream residues are distant from the PDZ surface, as observed in the connexin-43 peptide complex ^33^.

The interaction between the SARS-CoV-2-E-PBM peptide and the binding pocket of ZO-1 PDZ2 exemplifies a type II PBM-PDZ binding ^32^. Three hydrogen bonds are formed between the C-terminal carboxylate of Val at position 0 (Val0) of the PBM and the backbone amides of Tyr198, Gly199, and Leu200 of the “GLGF” loop in ZO1-PDZ2 (**Figure 1B**). A hydrogen bond is also formed between the backbone amide of Val0 and the carbonyl of Leu200 at the beginning of the β2 strand. Additionally, the carbonyl and the amide group of Leu at position -2 (Leu-2) interact through hydrogen bonds with the amide and carbonyl groups of Leu200, respectively. Also, the side chain of Leu−2 engages in a hydrophobic interaction, packing tightly against the side chain of Leu202 located in the β2 strand. An additional H-bond is observed between the carbonyl of Val-5 and the side-chain amide of Asn239 (**Figure 1B**). The structural insights into the ZO-1 PDZ2 domain in complex with the SARS-CoV-2 E C-terminal provide valuable clues for therapeutic targeting of this interaction.

### High-throughput HTRF screening uncovers 36 compounds targeting E/ZO-1 protein-protein interaction

To identify potential inhibitors, we employed a homogeneous time-resolved fluorescence resonance energy transfer (HTRF) approach to screen compounds that could disrupt the E/ZO-1 PPI. This variant of FRET uses a persistent fluorescence donor, enabling measurements after a defined delay following excitation.

For the assay, we used an N-terminal His-tagged construct encompassing the cytoplasmic region of the E protein (residues 45-75) and a GST-tagged construct of the ZO-1 PDZ2 domain. Various in vitro binding parameters were tested in 96-well low-volume plate, including incubation time and temperature, as well as the concentrations of purified proteins and antibodies, to determine the optimal conditions for complex formation. Our experimental optimizations ensured robust and reliable FRET efficiency, assessed using the HTRF ratio and Z1 factor to evaluate signal-to- background ratio ^34^. With 3 nM of GST-ZO1-PDZ2 incubated with 60 nM of His-E for 1 hour at room temperature, followed by an additional 1-hour incubation with labeled antibodies, we confirmed a strong signal indicative of complex formation. GST-ZO1-PDZ2 alone with buffer was used as negative control, whereas DMSO, used as an organic solvent for compound dissolution, did not impair PPIs or reduce the reaction system stability at its final concentration of 5%.

Using a subset of the CNRS Chimiothèque Nationale, a focused library of 1,000 putative PPI inhibitors solubilized in pure DMSO ^31^, we conducted a primary screening at a concentration of 20 µM per compound, tested in duplicate. Our screening achieved a Z1-factor above 0.5 for all the plates, confirming the robustness and high-quality performance of our assay. We identified 36 promising hits with a residual HTRF ratio (% residual HTRF ratio) below 30% (Table 2). Histograms illustrating the percentage of inhibition of the normalized HTRF ratio signal are presented in **Figure 2**.

**Figure 2.**
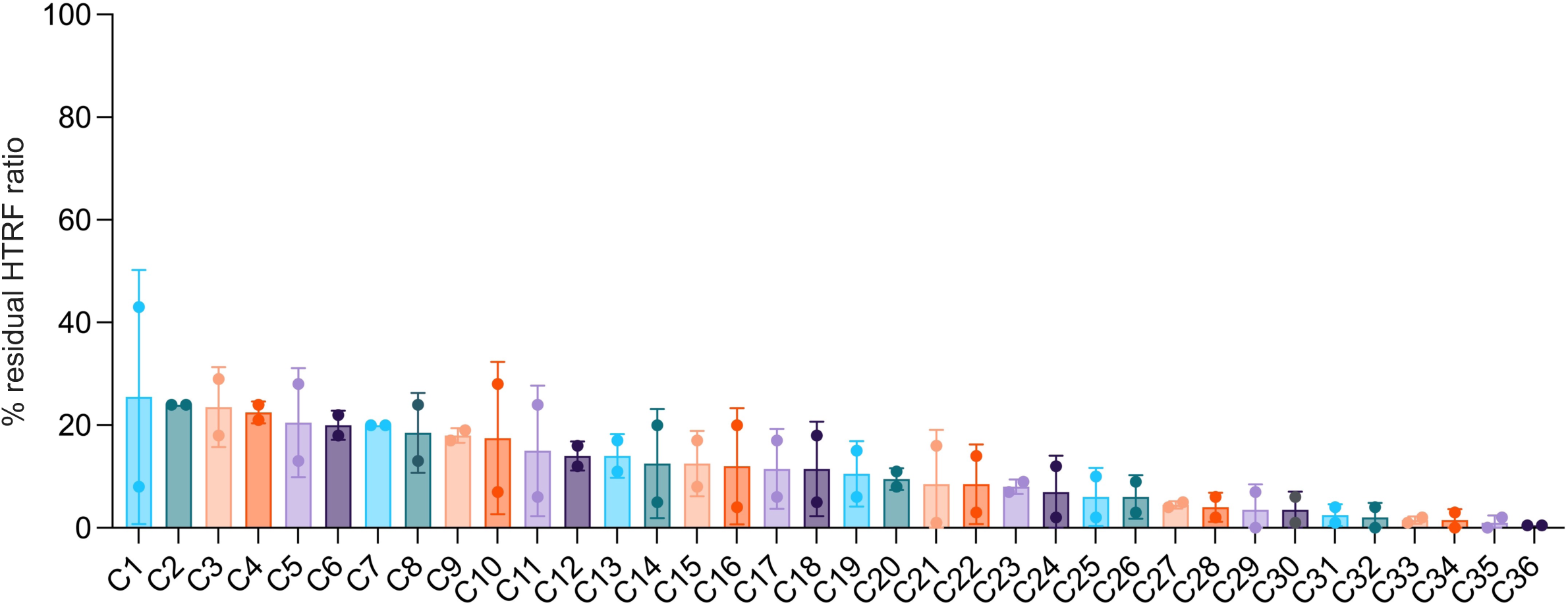
HTRF-based screening of compounds targeting the interaction between ZO-1 PDZ2 and the SARS-CoV-2 E protein PBM. The assay was conducted in 96-well, white, low-volume HTRF plates using 5 µL of purified GST-ZO-1-PDZ2 at a final concentration of 3 nM, 5 µL of purified His₆-Ecyto at a final concentration of 60 nM, and 1 µL of each of the 1,000 compounds from a focused-PPI chemical library. The histogram shows the % residual HTRF ratio. Among the 1,000 compounds screened, only the 36 compounds with % residual HTRF ratio below 30% were classified as primary hits (C1 to C36). Horizontal lines indicate the mean with SD (n=2).

**Table 2.** Activity and Toxicity Metrics of the 36 primary identified hits. This table summarizes the % residual HTRF ratio from the primary screening, along with IC₅₀ values determined from HTRF-based serial dilution assays and CC₅₀ values obtained from cytotoxicity dose-response assays. The CC₅₀/IC₅₀ ratio was calculated to determine the selectivity index (SI). For compounds that did not exhibit cytotoxicity, an SI threshold was estimated by using the highest tested concentration (20 µM) as the CC₅₀ limit. Dashes indicate data that were either not detected or not quantifiable. Values highlighted in red indicate compounds that fall outside the predefined selection thresholds, set at 25 µM for IC₅₀ (inhibitory potency) and 10 µM for CC₅₀ (cytotoxicity), and were therefore excluded from further consideration.

Upon further analysis of the screening results, we noted a spectrum of inhibitory effectiveness among the compounds. Compound C36, the most effective, exhibited a % residual HTRF ratio of just 0.5 ±0.0 %, highlighting its strong potency as an inhibitor of the E/ZO-1 interaction. Other notable compounds include C35 and C36, with residual % residual HTRF ratio of 1 and 0.5, respectively. The compounds demonstrated varying levels of potency to prevent the E/ZO-1 interaction, contributing to a diverse set of candidates for further investigation.

### Fourteen key inhibitors of the E/ZO-1 interaction emerge from the evaluation of cytotoxicity and dose-response evaluations

To evaluate the cytotoxicity of the compounds, we tested the 36 initial hits on Vero-E6 cells using a 9-point, two-fold serial dilution with inhibitor concentration ranging from 20 µM to 78 nM (**Supplementary Figure 2**, Table 2). Cell viability was measured using the XTT assay, allowing us to determine the CC₅₀ values of the compounds (Table 2). Compounds with a CC₅₀ lower than 10 µM were excluded from further analysis. Notably, compounds such as C16 (2.8 µM), C25 (5.7 µM), and C4 (9.1 µM) posed higher cytotoxicity risks and were discarded.

In parallel, we repeated the HTRF assays for all 36 compounds using a 10-point, two-fold serial dilution, with concentrations ranging from 100 µM to 195 nM (**Supplementary Figure 3**, Table 2). This approach aimed to assess both reproducibility and dose-dependent inhibitory effects, enabling calculation of IC₅₀ values (Table 2). Compounds with IC₅₀ values exceeding 25 µM were excluded based on our selection criteria. Among the inhibitors, C6 emerged as the most potent with an IC₅₀ of 0.4 µM, while C20 and C35 displayed low IC₅₀ values of 1.5 µM and 1.9 µM, respectively, combined with high CC₅₀ values (21.6 µM and 20.7 µM), indicating both strong inhibition and low cytotoxicity. C29 and C32 also had notable IC₅₀ values of 1.1 µM and 1.9 µM, alongside reasonable CC₅₀ thresholds, making them good candidates. Conversely, C22 or C33 exhibited elevated IC₅₀ values, placing them out of the preferred range for effective interaction inhibition.

Selectivity indices (SI) were calculated for all compounds, with several showing SI values greater than 1 and some exceeding 10, underscoring the identification of multiple selective candidates (Table 2). Based on cytotoxicity and dose-response inhibition assays, 14 promising compounds were selected, with HTRF IC₅₀ values ranging from 0.4 to 21.8 µM. The chemical structures of these selected compounds are shown in **Figure 3**. The compounds exhibit complex organic structures, predominantly characterized by fused aromatic rings and distinct functional groups. Many of these compounds incorporate fused aromatic or heteroaromatic rings, forming polycyclic structures. Several molecules possess functional groups such as hydroxyl (–OH), methoxy (–OCH₃), amide (–CO–NH–), and isocyanate (–NCO), which are characteristic of highly functionalized PPI compounds designed to promote interactions such as hydrogen bonding, thereby influencing their chemical and biological properties. Compounds such as C20, C32 and C35 exhibit structural symmetry, indicating a regular arrangement of functional groups and rings, characteristic of ditercalinium-type compounds with only slight variations in their substituents. Three compounds possess potentially reactive functional groups: compound C10 contains an isocyanate group, compound C19 features an aromatic nitro group, and compound C36 includes a catechol moiety with potential redox activity. Compound C29 presents a linear core structure flanked by fused rings at the ends, whereas other compounds, like C32, adopt a more complex, polycyclic configuration. Compound C28 is identified as a metal chelator, suggesting its ability to bind metal ions, a feature not common to all compounds. Among the most active compounds, compound 19 (C19) is constructed around a quinoline substituted in position 2 by a catechol masked in the form of methyl ether and in position 7 by a nitroaromatic which can be the origin of redox cycling reactions.

**Figure 3.**
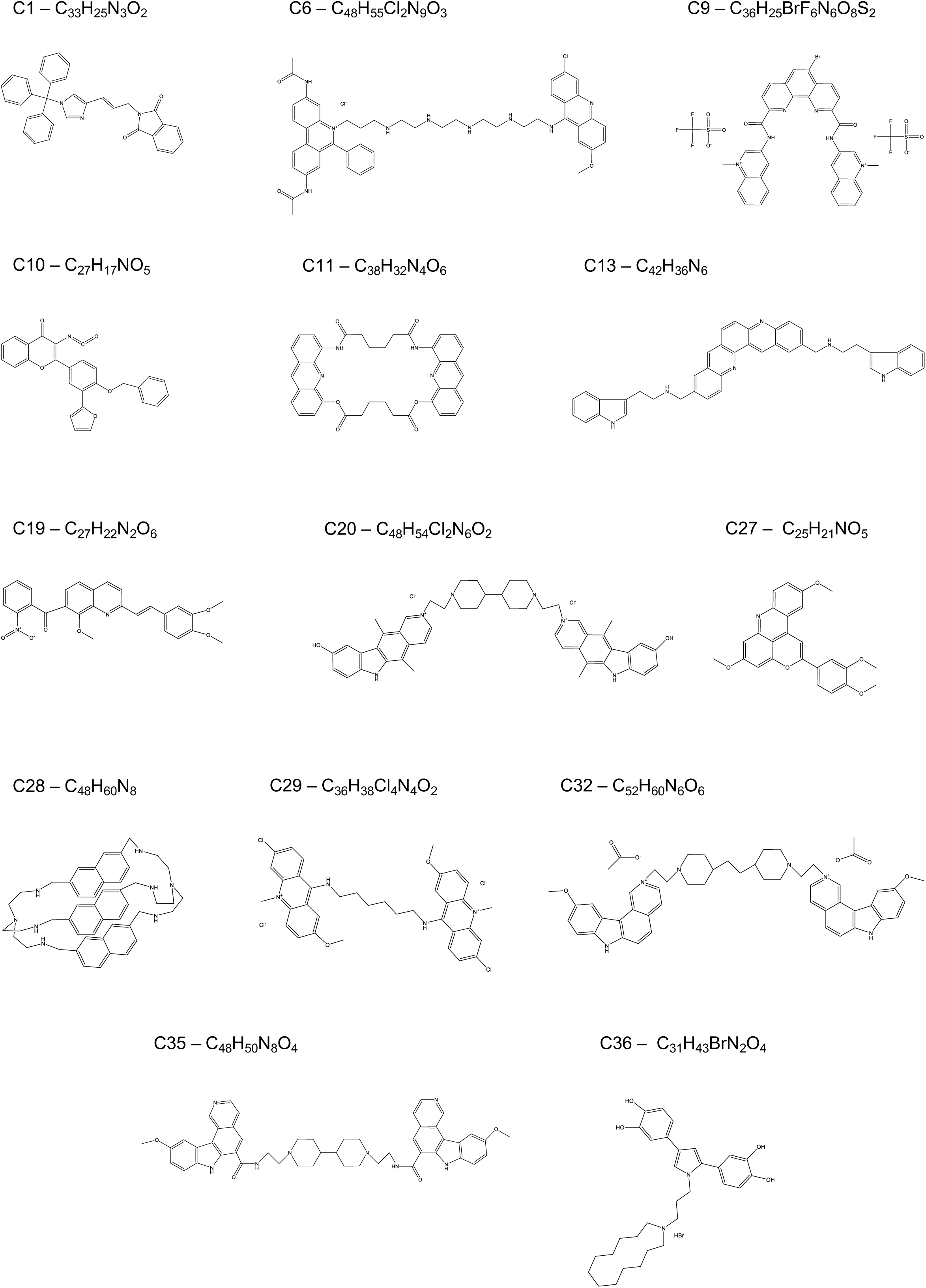
Chemical structures of the 14 selected compounds. The 14 compounds shown were selected based on results from cytotoxicity and inhibitory dose-response assays. All chemical structures were generated using ChemDraw software.

Due to their polycyclic structures and specific functional groups, the compounds may interfere with the interaction between the SARS-CoV-2-E-PBM peptide and the binding pocket of ZO-1 PDZ2. For example, fused aromatic rings and functional groups like amides or isocyanates could potentially bind to the PDZ2 domain of ZO-1, inhibiting its interaction with the E protein PBM. Additionally, the presence of positive charges in at least eight compounds with more than two charges may enhance electrostatic interactions with charged residues in the PDZ2 domain, thereby increasing their affinity for this target. In summary, despite the structural diversity among these compounds, their ability to interact with the PDZ2 domain of ZO-1 through π- stacking, electrostatic interactions, or metal ion coordination could render them effective in inhibiting the E protein PBM-ZO-1 PDZ2 interaction.

### Docking analysis identifies key binding sites for selected compounds on the ZO-1 PDZ2-E PBM complex

To further investigate the binding modes of the 14 selected compounds, we performed docking simulations using the DOCK6 program ^35^. As a template, we employed the X-ray structure of the ZO-1 PDZ2 domain alone (PDB ID: 2RCZ). All compounds were docked successfully, with the molecules adopting various orientations (**Figure 4**, **Supplementary Table 1**). The docking poses demonstrated a tendency to cluster in a central region at the interface of the swapped dimer, positioned between the two PDZ monomers. Moreover, among the top poses for the five compounds C1, C10, C19, C27 and C36, some were found in proximity to the PBM peptide-binding site close to the α2 helix (**Figure 4**). These findings suggest that while certain compounds may directly compete or interfere with the PBM for binding, some might also disrupt the interaction by targeting adjacent regions crucial for the structural stability of the dimer. The preferential localization of the compounds to be docked within this central pocket and near the PBM peptide-binding site highlights a conserved hotspot for ligand binding, which could be exploited for therapeutic targeting of the ZO-1 PDZ2/SARS-CoV-2 E interaction. Furthermore, control docking simulations on the structure of ZO-1 PDZ2 dimer bound to the PBM peptide that we obtained revealed no compounds located near the PBM-binding pocket (**Supplementary Figure 4**), consistent with a competitive inhibition mechanism.

**Figure 4.**
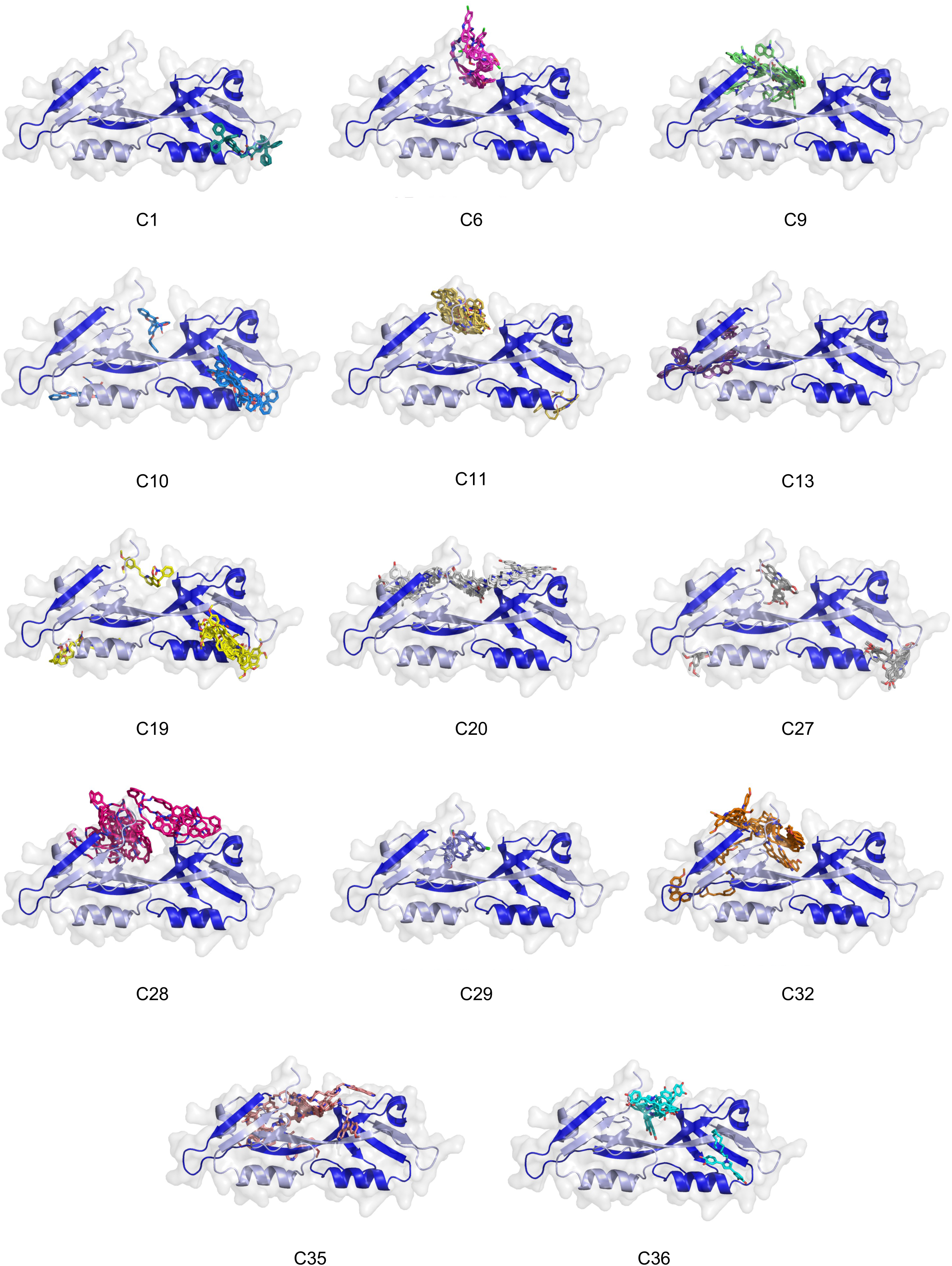
Docking poses of the top 14 compounds targeting the ZO-1 PDZ2 dimer. The figure shows the best binding poses of the 14 compounds targeting the swapped dimer of ZO-1 PDZ2 using DOCK6 docking simulations. The PDZ is depicted in cartoon. Each compound’s docking pose is represented, positioned within key binding regions of the ZO-1 PDZ2 dimer. The compounds are primarily located at the dimer interface or near the PBM binding site such as C10, C19 or C27.

### A standout molecule among the hits emerges as promising for its antiviral activity

To investigate the potential antiviral effects of the 14 selected compounds directly extracted from the library without further chemical optimization, we used a recombinant SARS-CoV-2 virus engineered to express NanoLuciferase as a reporter gene. This system allowed us to quantify viral replication by measuring luminescence from the NanoLuciferase secreted into the infected cell culture supernatant, which directly correlates with viral load (**Supplementary Figure 5**). Among the 14 tested compounds, C19 stood out by inducing a greater than two-log reduction in luminescence at concentrations above 10 µM, indicating a significant impact on viral replication.

This compound (**Figure 5A**), for which all-specific data are summarized in **Figure 5**, was predicted in 8 out of 9 top docking poses to occupy the PBM interaction pocket, suggesting a potential competitive binding mechanism (**Figure 5B**). Supporting this hypothesis, when docking was performed using our structure of the ZO-1 PDZ2 domain in complex with the E PBM peptide, all binding poses of C19 were redirected toward the interface between the two monomers, rather than the PBM-binding site (**Figure 5C**).

**Figure 5.**
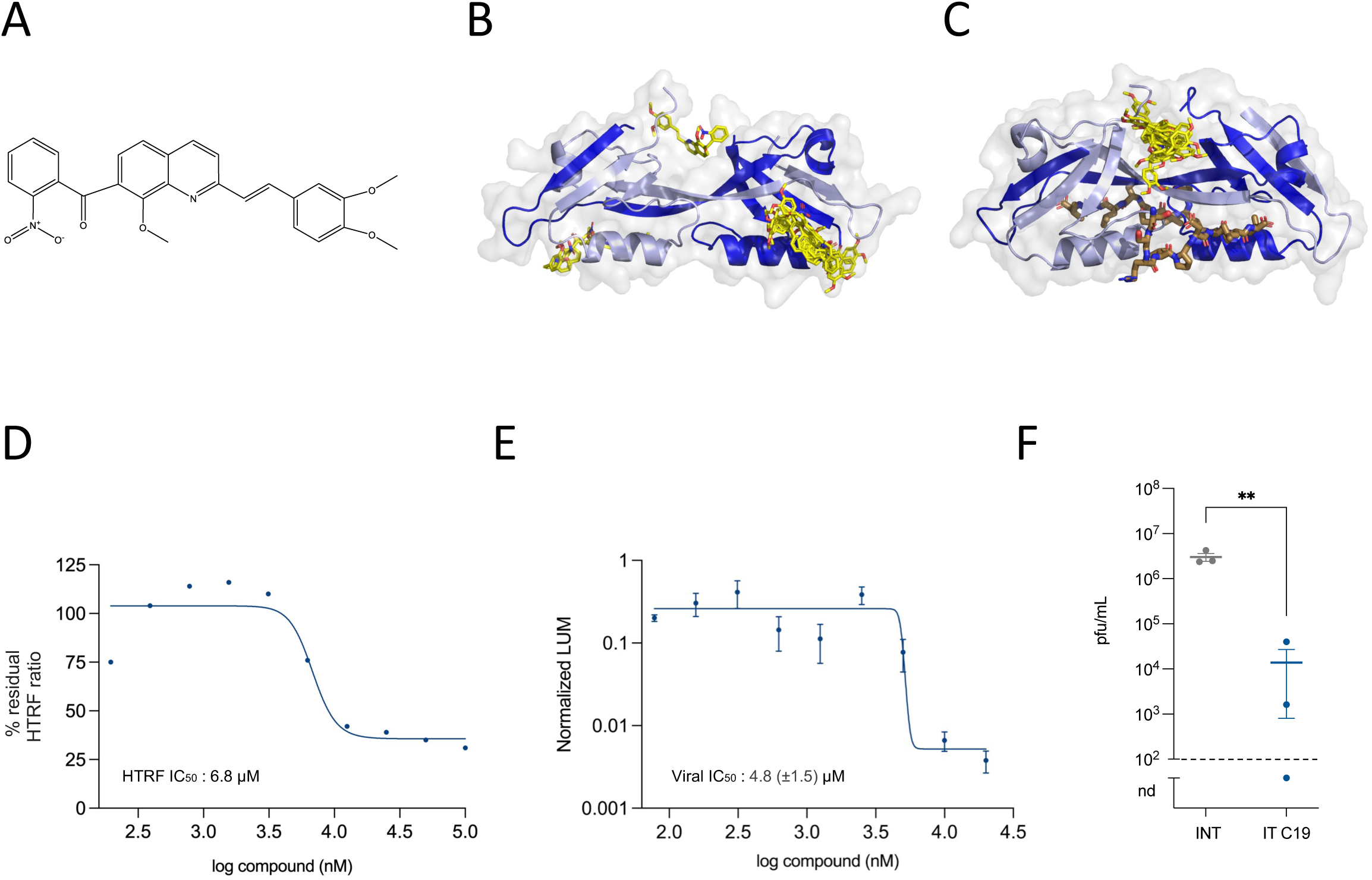
Characterization of C19, a promising antiviral compound. **A**. Chemical structure of C19, generated using ChemDraw software. B.C. Docking poses of C19 targeting the ZO-1 PDZ2 swapped dimer. Panels show the best predicted binding poses of C19 when docked to the ZO-1 PDZ2 dimer either in its free form (**B**) or in complex with the E protein PBM (**C**), using DOCK6 simulations. The PDZ domains are shown in cartoon representation. The compound is positioned within key binding regions of the ZO-1 PDZ2 dimer, preferentially occupying the PBM binding site. **D**. HTRF inhibition dose-response curve of C19. The assay was performed in 96-well, white, low-volume HTRF plates with 5 µL of purified GST-ZO-1-PDZ2 (final concentration: 3 nM), 5 µL of purified His₆-Ecyto (final concentration: 60 nM), and 1 µL of C19, tested in a 10-point, twofold serial dilution (100 µM to 195 nM). The resulting curve represents the percentage of residual interaction (%R) based on the normalized FRET signal. Sigmoidal curve was generated using nonlinear regression in Prism software, plotting C19 concentration against the normalized response (n=1). **E.F**. Antiviral activity of C19. The assay was conducted in 96-well culture plates using Vero-E6 cells (40,000 cells/well) infected with a NanoLuciferase-expressing recombinant SARS-CoV-2 virus at a MOI of 0.001. Cells were incubated for 72 hours in the presence of C19, applied as a 9-point, twofold serial dilution ranging from 20 µM to 78 nM. Viral replication was assessed by measuring luminescence in the cell culture supernatant using the Nano-Glo assay kit (**E**). Data were normalized to both infected untreated and uninfected untreated control conditions. Sigmoidal curve was generated using nonlinear regression in Prism software, plotting C19 concentration against the normalized luminescence signal (n=3). Viral quantification was performed using plaque assay on Vero-E6 cells (F). Plaques were observed at 3 days post-infection (dpi) following infection with serial dilutions of supernatants collected from infected non-treated (INT) or infected treated (IT) cells, treated for 72 hours with 20 µM of compound C19. Viral titers are reported as plaque-forming units per milliliter (PFU/mL). Horizontal lines represent the mean ± SEM, and dashed lines indicate the limit of detection (n=3; nd = not detected). Statistical significance was determined by t-test (p < 0.005).

Compound C19 exhibits an IC₅₀ of 6.8 µM for inhibition of the E/ZO-1 interaction (**Figure 5D**), as measured by HTRF, and an IC₅₀ of 4.8 ± 1.5 µM for inhibition of viral replication (**Figure 5E**), determined via luminescence assays in infected NanoLuciferase-expressing virus-infected cells. Viral titration by plaque lysis assay confirmed the antiviral effect, showing a viral titer more than 2 log units lower compared to untreated infected cells (**Figure 5F**).

## Discussion

The SARS-CoV-2 E protein plays a multifaceted role in viral pathogenesis. Various molecules and strategies have been investigated to inhibit the SARS-CoV-2 E protein, although research in this area is still ongoing ^12^. These include direct inhibitors, peptide antagonists, antibody-based approaches, repurposed drugs, and computationally designed molecules. As understanding of the E protein’s role in viral pathogenesis and its interactions with host proteins continues to evolve, new and more effective inhibitors are likely to emerge. While monoclonal antibodies and ion channel blockers have been developed, approaches targeting the PDZ-binding motif remain unexplored ^12^.

We previously identified ZO-1 PDZ2 as a binding partner of the SARS-CoV-2 E protein and demonstrated that its PBM preferentially binds to ZO-1 PDZ2 in vitro with a dissociation constant (Kd) of 15 μM ^15,24^. ZO-1 is a critical component of tight junctions, which are essential for maintaining epithelial and endothelial barrier integrity. During SARS-CoV-2 infection, ZO-1 has been implicated as a key host factor targeted by the viral E protein, particularly through its C-terminal PBM. We previously highlighted that this PBM acts as an important factor in pathogenesis, demonstrating how disruption of the ZO-1 network contributes to tight junction and epithelial barrier compromise ^22^. This breakdown of barrier integrity may play a significant role in the development of ARDS observed in severe COVID-19 cases.

The crystal structure of the ZO-1 PDZ2 domain in complex with the SARS-CoV-2 E PBM reveals key insights into the molecular determinants of the interaction and provides a foundation for the development of novel therapeutic strategies. The adoption of a swapped dimer conformation stabilized by the reciprocal β1/β6 and β2/β3 interactions provides a robust framework for peptide binding. The small RMSD between the structures of the PDZ in complex with the SARS-CoV-2 E PBM peptide and the connexin-43 peptide, which also harbors a PBM (DLEI_COOH_), further validates the conservation of the interaction mode. This emphasizes the structural similarity and adaptability of the ZO-1 PDZ2 domain in engaging diverse PBM ligands.

The high-throughput HTRF screening identified 36 compounds capable of disrupting the E protein-ZO-1 interaction, confirming the effectiveness of this method for discovering PPI inhibitors. Cytotoxicity and dose-response assessments reduced these down to 14 promising candidates. Docking simulations further revealed that some compounds preferentially bind within or near the PBM-binding groove, while others engage adjacent regions at the interface between the two PDZ monomers, suggesting multiple potential mechanisms for disrupting the interaction. Among these, compound C19 emerged as a standout due to its potent antiviral activity, demonstrated by a significant reduction in viral replication observed using a recombinant SARS-CoV-2 NanoLuciferase reporter system. Our study highlights C19 as a promising antiviral candidate that effectively disrupts the E protein-ZO-1 interaction. However, further optimization of this compound is necessary to enhance its specificity, potency, and pharmacokinetic properties. The structural insights gained from the ZO-1 PDZ2–E PBM complex and docking studies provide a framework for designing next-generation inhibitors targeting this critical host-virus interaction. Such inhibitors could not only mitigate the direct effects of SARS-CoV-2 infection but also provide a blueprint for addressing other pathogens that exploit similar host interactions.

Disrupting this interaction could preserve barrier integrity, reduce inflammation, and alleviate COVID-19 severity. Since ZO-1 is expressed in multiple organs, such as the gastrointestinal tract, kidneys, and brain ^29,36,37^, the E protein’s interaction with ZO-1 may contribute to systemic complications like gastrointestinal symptoms, acute kidney injury, and neurological issues observed in some COVID-19 patients. Additionally, the E/ZO-1 interaction may compromise the endothelium, leading to vascular complications, including blood clots, strokes and multiorgan failure ^38,39^. In conclusion, targeting this interaction may reduce viral fitness and help mitigate both pulmonary and systemic complications of COVID-19, underscoring the importance of continued efforts to develop inhibitors of key host–virus interactions and paving the way for novel antiviral strategies.

## Materials and Methods

### Expression constructs

The pGEX-2T plasmid encoding amino acids 186–262 of human ZO-1 was obtained from Addgene (59735, Addgene) ^32^. The construct features a thrombin cleavage site inserted between the GST tag and the N-terminal PDZ sequence. The cDNA sequence encoding amino acids 45-75 of the SARS-CoV-2 E protein was synthesized and then cloned into the pET-15b vector by GenScript. In this construct, the E protein sequence is appended with a N-terminal His tag, which is linked by a TEV protease cleavage site.

### Protein expression and purification

Vectors encoding His-E_cyto_ or GST-ZO-1-PDZ2 constructs were transformed into E. coli BL21 Star (DE3) cells (EC0114, Invitrogen™). The bacteria were cultured in LB medium supplemented with 100 mg/L ampicillin at 37°C. Protein expression was induced by adding 0.2 mM IPTG when the OD_600nm_ reached 0.8–1.0, and cultures were incubated overnight at 18°C. Following induction, the bacteria were harvested, and the PDZ2 domain of ZO-1 or Ecyto were resuspended in a lysis buffer composed of 50 mM Tris (pH 7.5), 250 mM NaCl, and 2 mM β-mercaptoethanol. The buffer was supplemented with one EDTA-free protease inhibitor tablet per 50 mL (COEDTAF-RO, Roche Diagnostics) and more than 250 units per liter of culture of benzonase (E1014-25KU, Sigma-Aldrich). The cells were lysed under high pressure using a CELLD homogenizer (1.3 kBar, 4°C). Cell debris was removed by centrifugation at 30,000 × g for 1 hour at 4°C. Proteins fused to GST tag were purified using a GSTrap HP affinity chromatography column (17528201, GE Healthcare), while proteins fused to His tag were isolated using nickel-chelated HiTrap Chelating HP columns (17524801, GE Healthcare). For crystallographic purposes, an additional step was applied to the GST-ZO-1-PDZ2 protein bound to the column. An overnight cleavage at 16°C was performed with Thrombin protease (SAE0147, Sigma-Aldrich) to remove the GST tag. Cleaved and uncleaved proteins were then processed through a final step of size-exclusion chromatography using a Sephacryl S-100 HR 16/60 column (17116501, GE Healthcare). The chromatography buffer consisted of 50 mM Tris (pH 7.5), 150 mM NaCl, 0.5 mM TCEP, supplemented with one EDTA 2% protease inhibitors tablet per 100 mL of buffer (CO-RO, Roche Diagnostics).

### Peptide synthesis

The acetylated peptides containing the C-terminal PBM sequence of E protein (12 residues long) of SARS-CoV-2 (_ac_NLNSSRVPDLLV_COOH_) were synthesized in solid phase using the Fmoc strategy (Proteogenix, Schiltigheim, France). The peptides were resuspended in pure water to prepare desired stock solutions.

### Crystallization, data collection, and structure determination

The ZO-1 PDZ2 – E PBM complexes for crystallization were generated by mixing ZO-1PDZ domains with SARS-CoV-2 E protein PBM peptide at a ratio of 1:2, ensuring that at least 92% of complexes were formed. Initial screening of crystallization conditions was carried out by the vapor diffusion method using a Mosquito^TM^ nanoliter-dispensing system (TTP Labtech) following the established protocols ^40^. Sitting drops were set up using 400 nL of a 1:1 mixture of each sample protein and crystallization solutions (672 different commercially available conditions) equilibrated against a 150 μL reservoir in multiwell plates (Greiner Bio-One). The crystallization plates were stored at 4 °C in a RockImager1000® (Formulatrix) automated imaging system to monitor crystal growth. The best crystals were obtained in crystallization conditions containing 20% PEG 3350 and 0.2M NH4I. Crystals were then flash cooled in liquid nitrogen using the condition of crystallization supplemented with 30% (V/V) of glycerol as cryoprotectant. X-ray diffraction data were collected on the beamlines Proxima-1 Synchrotron SOLEIL (St. Aubin, France). The data were processed by XDS ^41^, and the structures were solved by molecular replacement with PHASER ^42^ using the search atomic models PDB id 2RCZ for ZO-1-PDZ2 as template. The positions of the bound peptides were determined from a Fo–Fc difference electron density maps. Models were rebuilt using COOT ^43^, and refinement was done with phenix.refine of the PHENIX suite ^44^. The crystal parameters, data collection statistics, and final refinement statistics are shown in Table 1. The structure factors and coordinates have been deposited in the Protein Data Bank under accession code 9HM2. All structural figures were generated with the PyMOL Molecular Graphics System, Version (Schrödinger).

### Design of a PPI-focused screening library

The French national chemical library (Chimiothèque Nationale) was used to design a focused screening library toward PPI. A chemoinformatic study was carried out to select 1,004 PPI-like compounds. First, the electronic version of the 62,281 compounds of the “Chimiothèque Nationale” was submitted to VSPrep, a workflow dedicated to preparing molecules for virtual screening ^45,46^. During the preparation of chemical libraries, VSPrep implements several metrics based on the distribution of molecular properties to characterize specific subsets such as PPI-like^47^. It also labels PAINS compounds (Pan Assay Interference Compounds) ^48^ using substructure search methods implemented in RDKit, a sofware package dedicated to chemoinformatics. 3 350 compounds were identified as PPI-like of which 117 labelled PAINS were removed, yielding 3223 compounds. The structural similarity of the compounds was assessed using FCFP6 fingerprints ^49^, calculated with RDKit. The compounds were then grouped according to their structural similarity using Butina algorithm ^50^. A Tanimoto coefficient threshold of 0.46 yielded 1,004 clusters from whose centroids were selected to build a library of 1,000 compounds that were solubilized in pure DMSO.

### HTRF binding assay between GST-ZO-1-PDZ2 and His6-Ecyto

The HTRF assay was conducted in 96-well, white, low-volume HTRF plate (66PL96025, Revvity) with a final reaction volume of 20 µL per well. The anti-GST antibody labeled with Terbium (Tb) cryptate as a donor (61GSTTLA, Revvity) was used to detect the GST-ZO-1-PDZ2 protein, while the anti-His6 antibody labeled with the d2 acceptor fluorophore (61HISDLA, Revvity) was employed to detect the His6-Ecyto protein. Experimental conditions were optimized to achieve the maximum HTRF ratio signal. 5 µL of purified GST-ZO-1-PDZ2 at a final concentration of 3 nM and 5 µL of purified His6-Ecyto at a final concentration of 60 nM, both diluted in assay buffer (50 mM Tris-HCl pH 7.5, 150 mM NaCl, 0.5 mM TCEP, 0.05% Tween-20, 1% BSA and one EDTA-free protease inhibitor tablet per 100 mL of buffer), were dispensed into the plate and incubated at room temperature for 1 hour. For the chemical library screening, 1 µL of compound (2 mM in DMSO) was spiked into dry wells. For each plate, columns 1 and 12 contained only 1 µL of DMSO to act as controls. Then, 4.5 µL of anti-GST mAb Tb-conjugate and 4.5 µL of anti-His6 mAb d2- conjugate, both diluted to a final concentration of 1X according to the manufacturer’s protocol, were added to the plate. The mixture was incubated for an additional 1 hour at room temperature. The fluorescence signal was measured with an Infinite M1000 Pro plate reader (TECAN) with the following settings: excitation at 340 nm, emission at 665 nm (for acceptor) and 620 nm (for donor) with a delay of 60 µs. The HTRF ratio is calculated for each well using the following formula: Signal at 665 nm/ Signal at 620 nm × 10^4^. and the Z’-factor^34^ was calculated to assess the robustness of the test, using the formula: 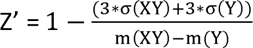 where “(XY)” refers to the mean (m) or standard deviation (σ) of the signal from wells containing both E and ZO-1 (positive controls), and “(Y)” refers to the mean or standard deviation of the signal from wells containing only ZO-1 (negative controls). The data from a plate are further analyzed when the Z’- factor is greater than 0.5. For each compound, the percentage of residual HTRF ratio was calculated via the following equation: % residual HTRF ratio = 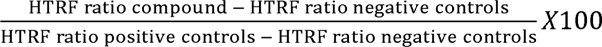. Compounds with a value of the % residual HTRF ratio below 30 were defined as primary hits

### IC_50_ HTRF determination

The primary hits were validated using HTRF under the same conditions described for the initial compound screening. Purified proteins were added to a 96-well plate containing 1 µL of each selected compound, diluted in DMSO and tested in a 10-point, 2-fold serial dilution ranging from a final well concentration of 100 µM to 195 nM, to determine the IC₅₀ values. For each compound, the interaction percentage, referred to as the HTRF residual interaction percentage, was calculated using the formula: % residual HTRF ratio = 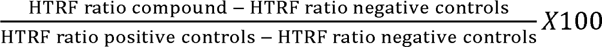. IC_50_ values were calculated from the experimental data (% residual HTRF ratio) using Prism Graphpad software and compounds with IC₅₀ values below 25 µM were classified as secondary hits.

### Cell viability assay

Cell viability was assessed using the CyQUANT™ XTT Cell Viability Assay (X12223, Invitrogen™). Vero-E6 cells were seeded into 96-well plates at a density of 4 × 10⁴ cells per well and incubated overnight. The following day, compounds were added to the cells in a 9-point, 2-fold serial dilution, ranging from a final concentration in well of 20 µM down to 78 nM, to determine the CC₅₀. Plates were cultured for 72 hours. To assess cell viability, the CyQUANT™ XTT Cell Viability Kit was used according to the manufacturer’s protocol. Briefly, the XTT reagent was prepared by mixing components A and B, and 70 µL of the prepared reagent was added to each well containing 100 µL of culture medium. After 4 hours of incubation at 37°C, absorbance was measured using an Infinit M1000 pro plate reader (TECAN) at 450 nm and 660 nm. Negative controls consisted of culture medium only, while positive controls involved a 1-hour treatment with 5% Triton X-100 diluted in culture medium to obtain 100% cell lysis. These controls enabled the calculation of the percentage of cell viability as a function of compound concentration and the subsequent determination of the CC₅₀. Compounds with CC₅₀ values above 10 µM were classified as secondary hits.

### Docking Simulations

We performed blind molecular docking simulations using the UCSF DOCK software (https://dock.compbio.ucsf.edu/DOCK_6/). The docking studies were conducted using the x-ray structure of the ZO-1 PDZ2 domain (PDB ID 2RCZ). The receptor structure included two PDZ2 domains of ZO-1 adopting a swapped dimer conformation. The receptor was prepared in MOL2 format with Chimera (removal of solvent molecules, addition of hydrogens and charges). The grid was generated to encompass all spheres detected with sphgen at the molecular surface of the receptor. The structures of the 14 selected compounds as ligands were prepared in MOL2 format with OpenBabel (addition of hydrogens, charges and generation of 3D conformation). Docking was carried out as flexible with DOCK6.9 using the grid-based score (van der Waals and electrostatic interactions). For each compound, at most 10 poses were ranked after clustering (2 Å RMSD threshold). The docking results were analyzed to identify potential binding sites and interactions between the compounds and the ZO-1 PDZ2 swapped dimer. Visualization of the docking poses and the analysis of intermolecular interactions were performed using PyMOL (Schrödinger, LLC). As a control, the same docking procedure was applied using the structure of the ZO1-PDZ2/E-PBM complex (PDB ID 9HM2) as receptor.

### Generation of the complete genome of Wuhan Nanoluciferase virus

Recombinant viruses expressing Nanoluciferase (nLuc) as a reporter gene were obtained by reverse genetics as previously described ^53^. Wuhan_nLuc was derived from the clone synSARS-CoV-2-GFP-P2A-ORF7a (clone #41) generated as described where the GFP reporter gene was replaced by the nLuc one. Briefly, the plasmid was linearized by EagI-HF (R3505, NEB) digestion, purified by a phenol chloroform process and transcribed in vitro into RNA using the T7 RiboMaxTM Large Scale RNA Production System (P1300, Promega) and the m7G(5’)ppp(5’)G RNA Cap Structure Analog (#S1411, NEB) kits. The synthesized RNA was finally purified using a classical phenol chloroform method and precipitated. Twelve μg of complete viral mRNA and 4 μg of plasmid encoding the viral nucleoprotein gene were electroporated on 81×110^6^ Vero-E6 cells resuspended in 0.8 mL of Electroporation Solution (MIR50114, Mirus Bio™ Ingenio™) using the Gene Pulser Xcell Electroporation System (1652660, Biorad) with a pulse of 270 V and 950 μF. The cells were then transferred to a T75 culture flask with 15 mL of DMEM supplemented with 2% FCS (v/v). The electroporated cells were incubated at 37 °C, 5% CO_2_ for several days until the cytopathic effect was observed. Then, the supernatant was harvested, aliquoted and frozen at −80 °C until titration. The viral sequence was controlled by Next Generation Sequencing.

### NanoLuciferase viral inhibition

To assess the impact of hit compounds on viral fitness, we used a recombinant virus expressing nLuc as a reporter gene. One day prior to infection, 40,000 Vero-E6 cells per well were seeded in a 96-well plate and incubated at 37°C with 5% CO₂ in 100 µL of medium supplemented with 5% fetal bovine serum (FBS). The following day, the cell culture medium was removed, and the cells were infected at a multiplicity of infection (MOI) of 0.001. After a 1-hour incubation period, the viral inoculum was removed. Hit compounds dissolved in 100% DMSO were diluted in cell culture medium without FBS, using a 9-point, 2-fold serial dilution, with final concentrations in wells ranging from 20 µM to 39 nM. Cells were cultured for 72 hours post-infection, after which luminescence was measured following the manufacturer’s protocol (N1120, Promega). Briefly, 10 µL of cell culture supernatant was diluted 10-fold into a final reaction volume of 100 µL in a flat-bottom white polystyrene 96-well plate (MSSWNFX40, Millipore®). The mixture was incubated for 10 minutes with the reconstituted reagent. Luminescence measurements were taken using a VICTOR® Nivo™ Plate Reader (PerkinElmer). Data are expressed as normalized luminescence using untreated infected cells and non-infected cells as controls.

### Lysis plaque viral titration

Viruses’ titers were determined by lysis plaque assay. 1.110^6^ Vero-E6 cells were seeded to each well of 6-well plates and cultured overnight at 37 °C, 5% CO_2_. Viruses were serially diluted in DMEM without FCS and 4001µl diluted viruses were transferred onto the monolayers. The viruses were incubated with the cells at 37 °C with 5% CO_2_ for 11hour. After the incubation, inoculum was removed, and an overlay medium was added to the infected cells per well. The overlay medium per well contained MEM 1X (prepared from MEM 10X (21430020, Gibco™), distilled water (15230162, Gibco™), L-glutamine (25030123, Gibco™) and gentamicin 10mg/ml (15710064, Gibco™)), 0.25% sodium phosphate (25080094, Gibco™), and 1X AVICEL (RC-581, Dupont). After 3 days of incubation, the plates were stained with crystal violet (V5265, Merck) and lyses plaques were counted to assess virus titer.

## Acknowledgements

This work was supported by: Institut Pasteur TASK FORCE COVID19, Institut Pasteur’s Programme Fédérateur de Recherche 5 (PFR5-Functional Genomics of the Viral Cycle funded by Fondation d’entreprise Michelin donation). Flavio Alvarez is recipient of a fellowship from the Institut Pasteur’s Programme Fédérateur de Recherche 5 (PFR-5 - Functional Genomics of the Viral Cycle). The production of recombinant viruses was performed thanks to the techniques developed by the Institut Pasteur’s Programme Fédérateur de Recherche 1 (PFR-1 - Reverse Genetics). We thank the staff of the crystallography platform at Institut Pasteur for carrying out robot-driven crystallization screening. We acknowledge synchrotron SOLEIL (Saint-Aubin, France) for granting access to their facility and the staff of Proxima1 for helpful assistance during the data collection. The authors acknowledge the French National Chemical Library (http://chembiofrance.org) for providing the compounds.

## Contributions

FA, NW and CCS conceived the project. FA and CCS supervised the project. FA, HML and CCS designed the experiments. SB provided the Chemical Library. FA, FL, QG, AH, BB, BSL, BF and CCS performed the experiments. FA, AM, HML, RADS and CCS analyzed the data. FA and CCS wrote the manuscript. All authors have read and edited the manuscript.

---

**Supplementary Figure 1.**
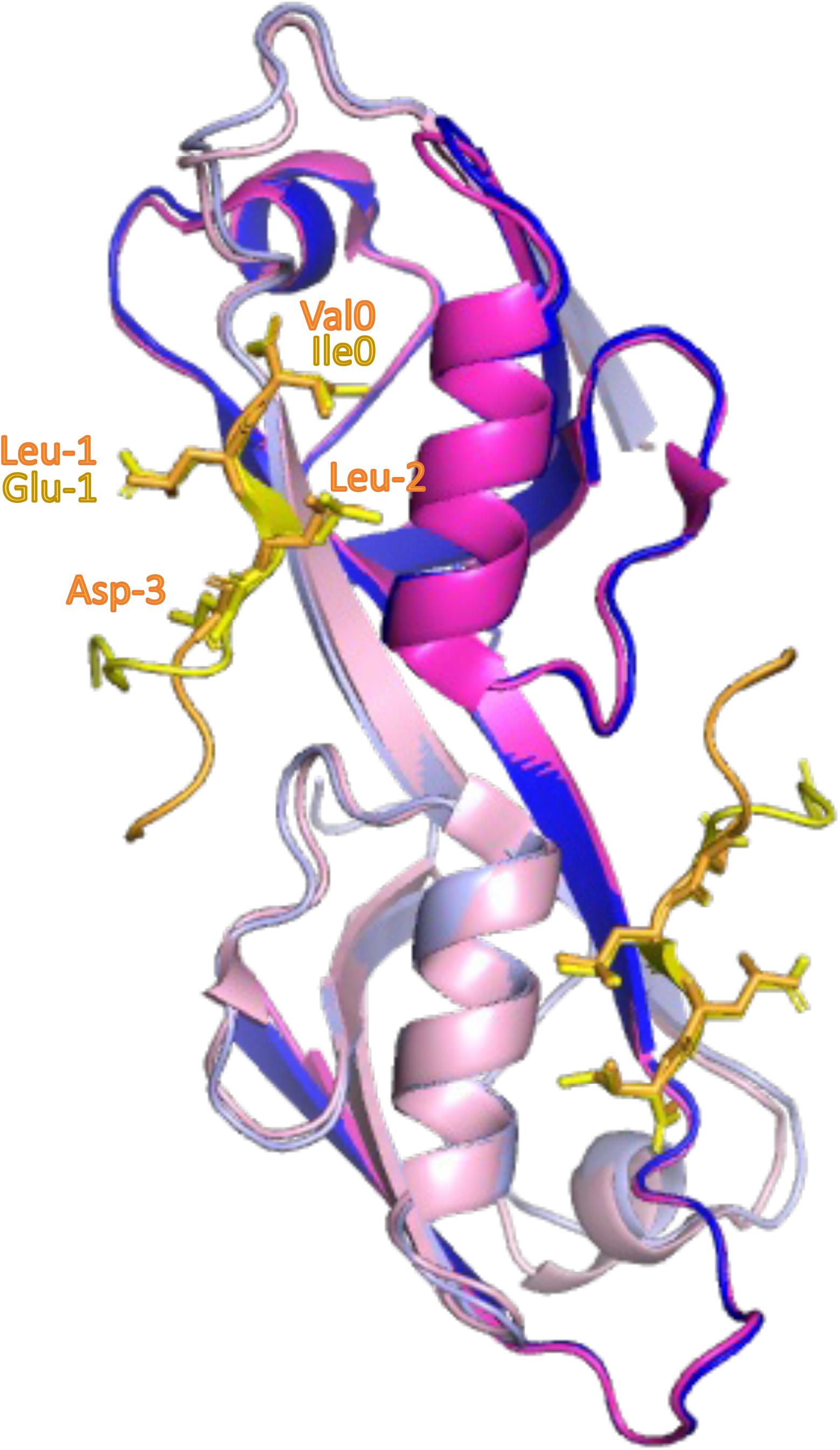
Overlay of the X-ray crystal structures of ZO-1 PDZ2 domain bound to the SARS-CoV-2 E PBM and connexin 43 PBM. Ribbon diagram of the final crystal structure model of the ZO-1 PDZ2 domain in complex with the SARS-CoV-2 E protein PBM peptide with one monomer colored in light blue and the other in dark blue. The bound peptides are shown in orange. The structure of the complex ZO-1 PDZ2 / Connexin 43 PBM peptide is superimposed with one monomer colored in pink and the other in magenta. The bound peptides are shown in yellow. The key residues of the PBM motifs are shown as sticks and labeled.

**Supplementary Figure 2.**
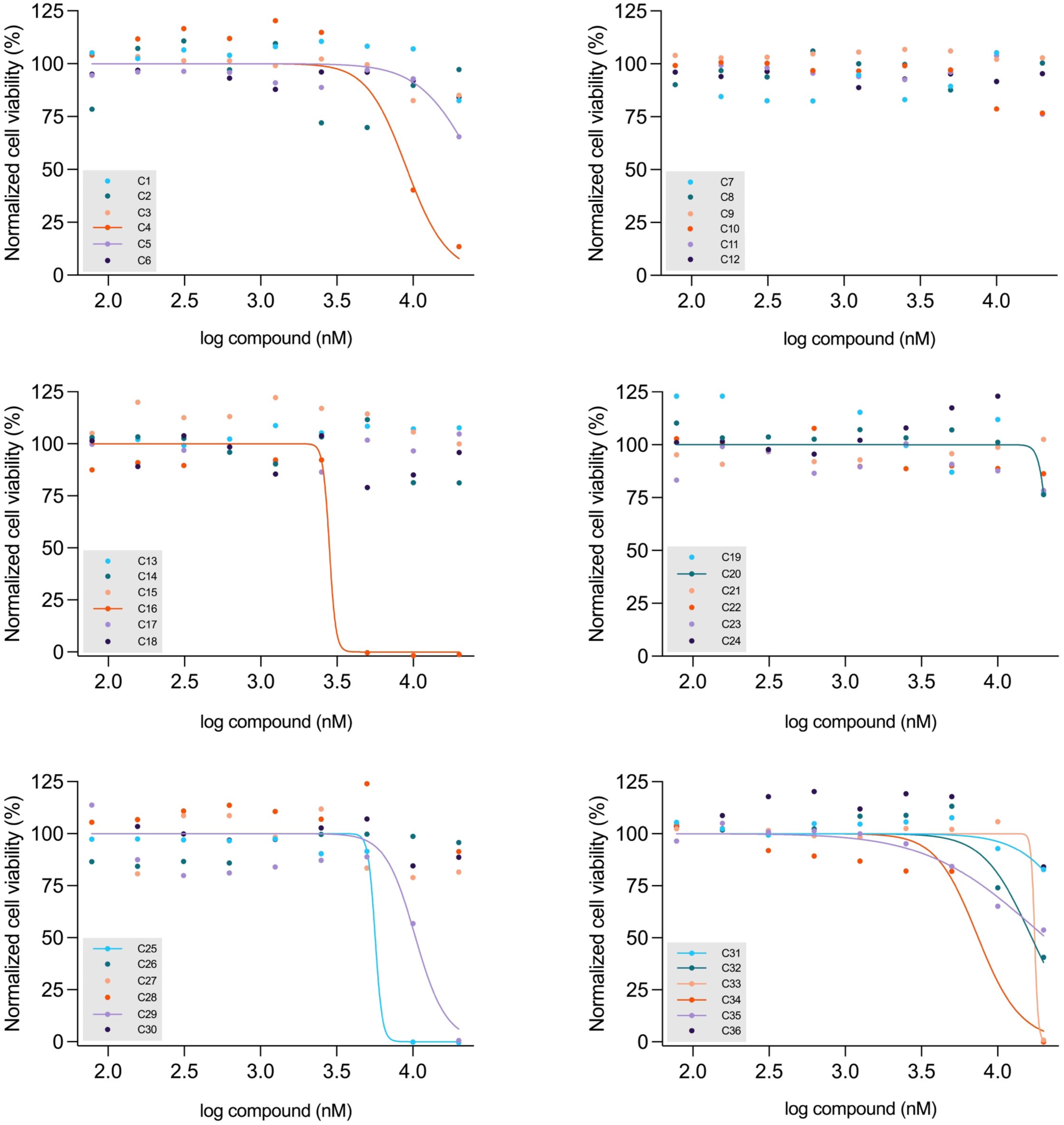
Cytotoxicity dose-response curves of the 36 primary identified hits targeting SARS-CoV-2 E protein interaction with ZO-1 PDZ2. The assay was performed in 96-well culture plates, with 40,000 Vero-E6 cells incubated for 72 hours in the presence of a 9-point, twofold serial dilution of each compound (concentrations ranging from 20 µM to 78 nM). Cell viability percentage was assessed using the XTT cell viability assay. Sigmoid curves were generated using nonlinear regression in Prism software, plotting inhibitor concentration against the normalized response (n=1). Related to Table 2.

**Supplementary Figure 3.**
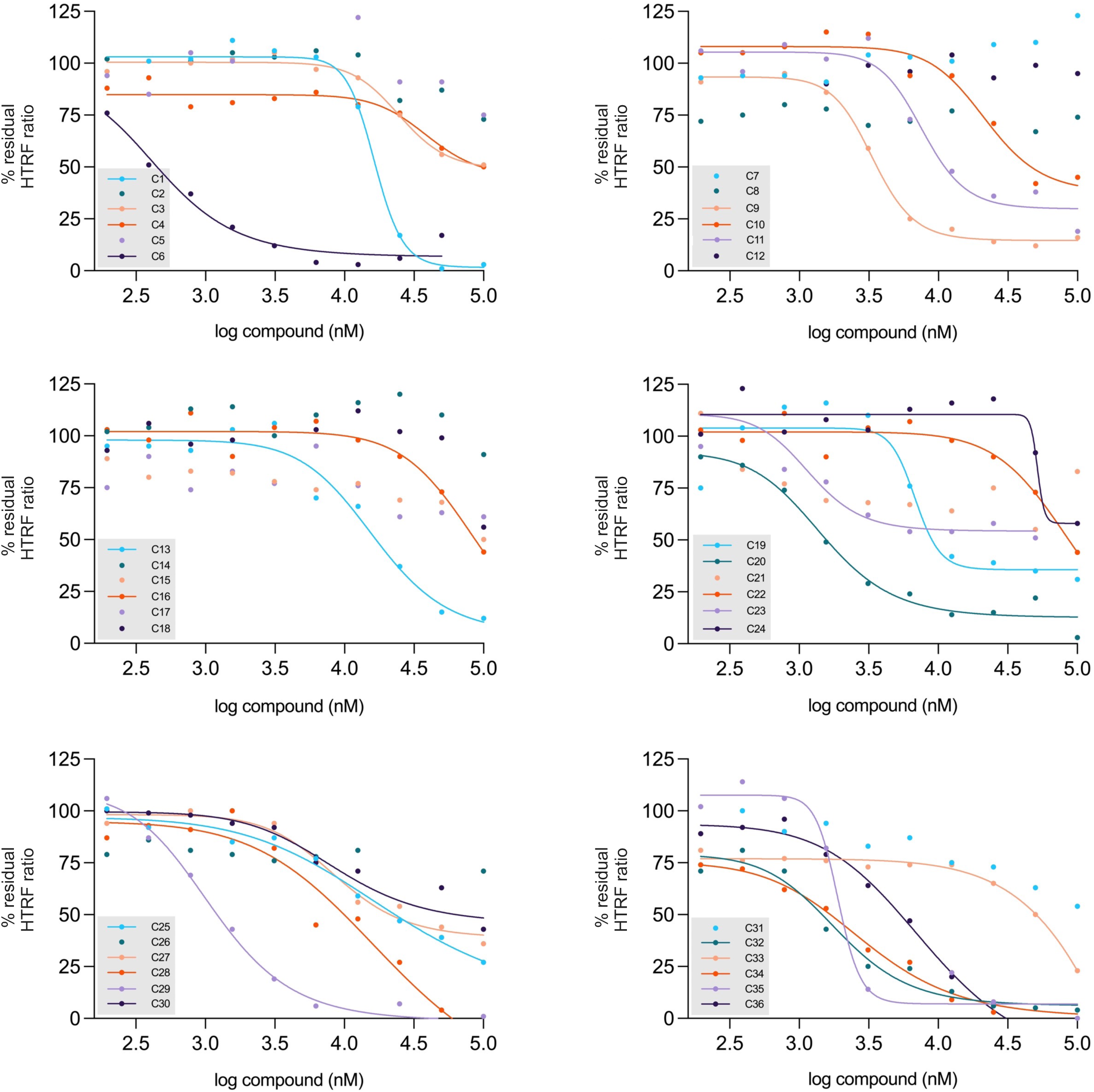
HTRF inhibition dose-response curves of the 36 primary identified hits targeting SARS-CoV-2 E protein interaction with ZO-1 PDZ2. The assay was conducted in 96- well, white, low-volume HTRF plates using 5 µL of purified GST-ZO-1-PDZ2 at a final concentration of 3 nM, 5 µL of purified His₆-Ecyto at a final concentration of 60 nM, and 1 µL of each of the 36 compounds identified in the primary HTRF screen. Compounds were tested in a 10-point, twofold serial dilution ranging from 100 µM to 195 nM. The resulting curves correspond to the % residual HTRF ratio as a function of the compound concentration. Sigmoidal curves were generated using nonlinear regression in Prism software, plotting inhibitor concentration against the normalized response (n=1). Related to Table 2.

**Supplementary Figure 4.**
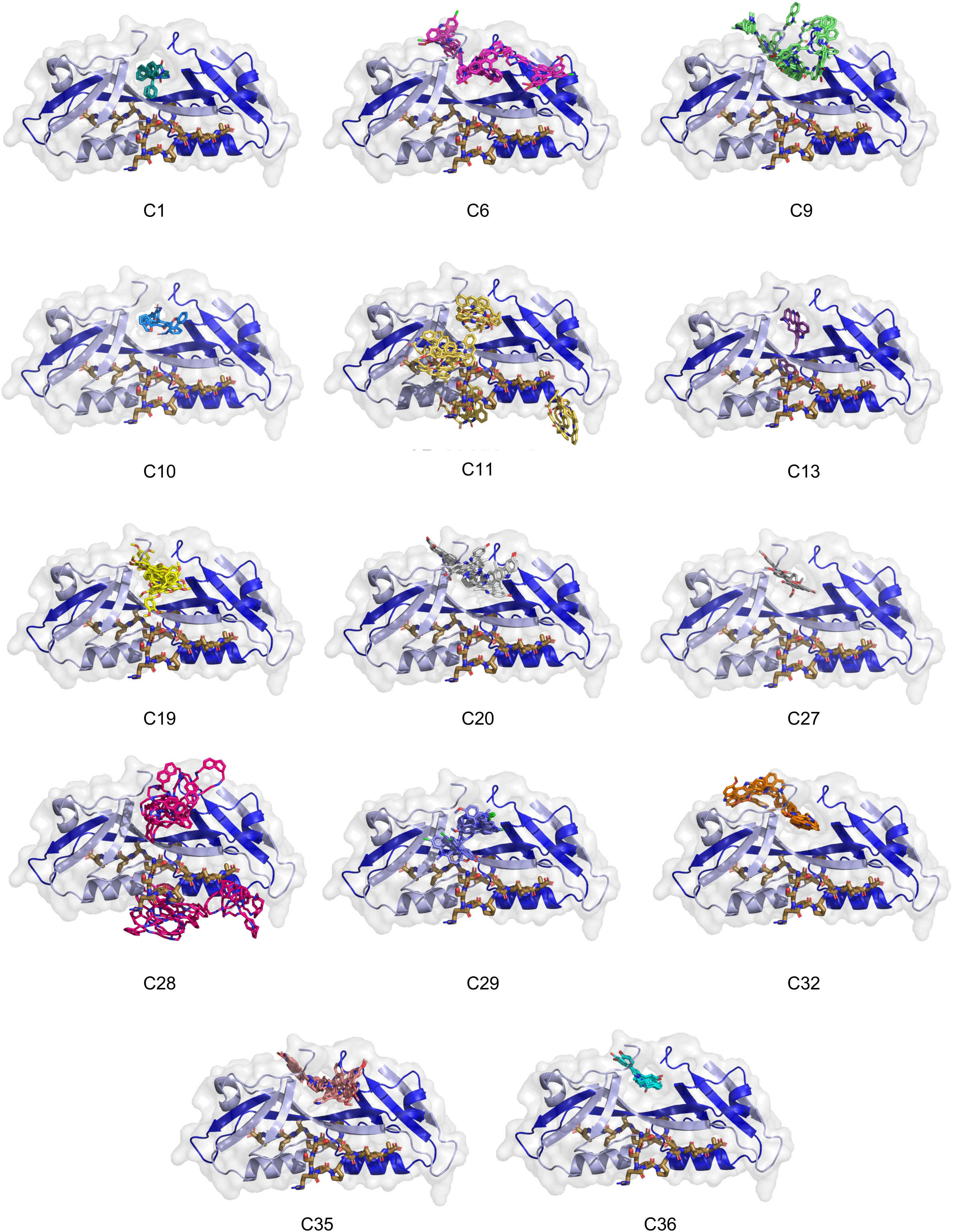
Docking control poses of the top 14 compounds targeting the ZO-1 PDZ2 swapped dimer in complex with the E protein PBM. Best poses of the 14 selected compounds docked into the ZO-1 PDZ2 swapped dimer using DOCK6 simulations in the presence of the E protein PBM. The PDZ domain is shown in cartoon representation, with the bound peptide colored in brown. Each compound’s docking pose is shown and can be compared to its respective pose obtained from docking to the free (unbound) swapped PDZ domains, illustrating the steric hindrance introduced by the peptide within the binding pocket. Related to Figure 4.

**Supplementary Table 1.** Docking parameters. Number of clustered poses and range of grid- scores of clustered poses for docking results using DOCK6 on ZO-1 PDZ2 dimer structure alone or in complex with E peptide. Related to Figure 4 and Supplementary Figure 4.

**Supplementary Figure 5.**
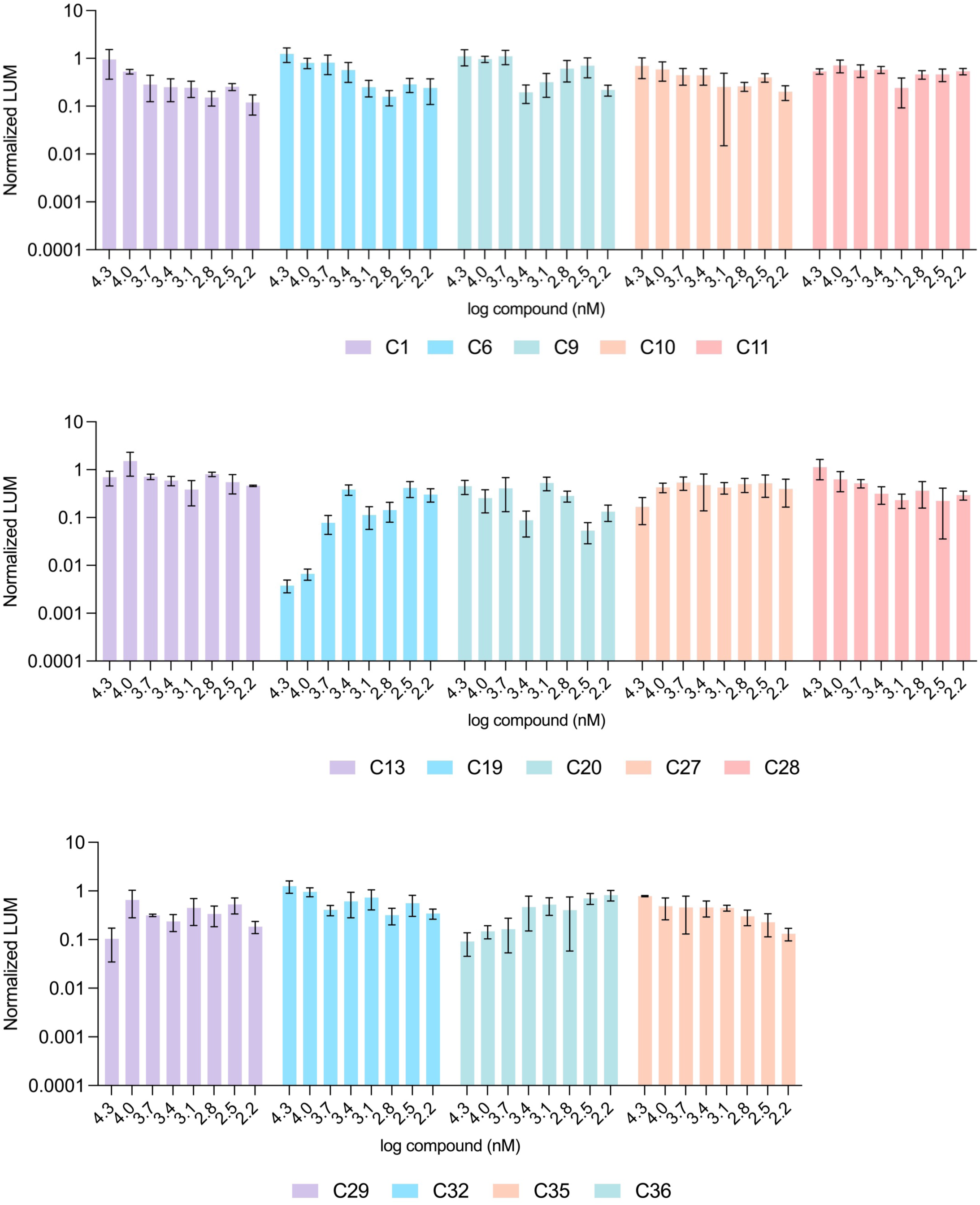
Antiviral activity of the 14 secondary screening hits. The assay was conducted in 96-well culture plates using Vero-E6 cells (40,000 cells/well) infected with a NanoLuciferase-expressing recombinant SARS-CoV-2 virus at a MOI of 0.001. Cells were incubated for 72 hours in the presence of each compound, applied as a 9-point, twofold serial dilution ranging from 20 µM to 78 nM. Viral replication was assessed by measuring luminescence in the cell culture supernatant using the Nano-Glo assay kit. To generate histograms, data were normalized to both infected untreated and uninfected untreated control conditions (n=3). Only C19 compound exhibits an antiviral effect, recorded at the highest tested concentrations. Related to Figure 5.

**ZO-1 PDZ2 / SARS-CoV-2-E-PBM peptide complex**
|  |  |
| --- | --- |
| **Wavelength** | 0.9786 |
| **Resolution range** | 54.120 - 1.721 (1.918 - 1.721) |
| **Space group** | P 21 21 21 |
| **Unit cell** | 30.127 76.157 76.922 90 90 90 |
| **Total reflections** | 180485 (7430) |
| **Unique reflections** | 13687 (684) |
| **Multiplicity** | 13.2 (10.9) |
| **Completeness (%)** | 52.9 (99.7) |
| **Mean I/sigma(I)** | 11.6 (1.7) |
| **Wilson B-factor** | 30.54 |
| **R-merge** | 0.121 (1.612) |
| **R-meas** | 0.126 (1.686) |
| **R-pim** | 0.034 (0.476) |
| **CC1/2** | 0.998 (0.617) |
| **Reflections used in refinement** | 13183 (77) |
| **Reflections used for R-free** | 684 (5) |
| **R-work** | 0.2030 (0.4132) |
| **R-free** | 0.2727 (0.5860) |
| **Number of non-hydrogen atoms** | 1492 |
| macromolecules | 1396 |
| ligands | 5 |
| solvent | 91 |
| **Protein residues** | 183 |
| **RMS(bonds)** | 0.006 |
| **RMS(angles)** | 0.82 |
| **Ramachandran favored (%)** | 99.43 |
| **Ramachandran allowed (%)** | 0.57 |
| **Ramachandran outliers (%)** | 0.00 |
| **Rotamer outliers (%)** | 0.64 |
| **Clashscore** | 2.83 |
| **Average B-factor** | 44.02 |
| macromolecules | 44.00 |
| ligands | 36.16 |
| solvent | 44.82 |

| Compound | HTRF residual | | Cytotoxicity CC<sub>50</sub><br>(μM) (n=1) | Selectivity<br>Index |
| --- | --- | --- | --- | --- |
| | Primary screening<br>percentage residual<br>ratio (%) (n=2) | IC50 (μM) (n=1) | | |
| C1 | 25.5 (± 24.7) | 16.4 | — | > 1.2 |
| C2 | 24 (± 0) | — | — | — |
| C3 | 23.5 (± 7.8) | 25 | — | > 0.8 |
| C4 | 22.5 (± 2.1) | 39.4 | 9.1 | 0.2 |
| C5 | 20.5 (± 10.6) | — | 26.5 | — |
| C6 | 20 (± 2.8) | 0.4 | — | >50 |
| C7 | 20 (± 0) | — | — | — |
| C8 | 18.5 (± 7.8) | — | — | — |
| C9 | 18 (± 1.4) | 3.4 | — | > 5.9 |
| C10 | 17.5 (± 14.8) | 21.8 | — | > 0.9 |
| C11 | 15 (± 12.7) | 7.7 | — | > 2.6 |
| C12 | 14 (± 2.8) | — | — | — |
| C13 | 14 (± 4.2) | 16.3 | — | > 1.2 |
| C14 | 12.5 (± 10.6) | — | — | — |
| C15 | 12.5 (± 6.4) | — | — | — |
| C16 | 12 (± 11.3) | 6.6 | 2.8 | 0.4 |
| C17 | 11.5 (± 7.8) | — | — | — |
| C18 | 11.5 (± 9.2) | — | — | — |
| C19 | 10.5 (± 6.4) | 6.8 | — | > 2.9 |
| C20 | 9.5 (± 2.1) | 1.5 | 21.6 | 14.4 |
| C21 | 8.5 (± 10.6) | — | — | — |
| C22 | 8.5 (± 7.8) | 87.2 | — | 0.2 |
| C23 | 8 (± 1.4) | — | — | — |
| C24 | 7 (± 7.1) | 51.6 | — | > 0.38 |
| C25 | 6 (± 5.7) | 21 | 5.7 | 0.27 |
| C26 | 6 (± 4.2) | — | — | — |
| C27 | 4.5 (± 0.7) | 8.7 | — | > 2.3 |
| C28 | 4 (± 2.8) | 20.3 | — | > 1 |
| C29 | 3.5 (± 4.9) | 1.1 | 10.4 | 9.45 |
| C30 | 3.5 (± 3.5) | — | — | — |
| C31 | 2.5 (± 2.1) | — | 38 | — |
| C32 | 2 (± 2.8) | 1.9 | 16.7 | 8.8 |
| C33 | 1.5 (± 0.7) | 483.1 | 17.5 | 0.04 |
| C34 | 1.5 (± 2.1) | 2.9 | 7.4 | 2.6 |
| C35 | 1 (± 1.4) | 1.9 | 20.7 | 10.9 |
| C36 | 0.5 (± 0) | 8.1 | — | > 2.5 |

## References

1. Carabelli, A. M. et al. SARS-CoV-2 variant biology: immune escape, transmission and fitness. Nat. Rev. Microbiol. 21, 162–177 (2023).

2. Markov, P. V. et al. The evolution of SARS-CoV-2. Nat. Rev. Microbiol. 21, 361–379 (2023).

3. Huang, C. et al. Clinical features of patients infected with 2019 novel coronavirus in Wuhan, China. Lancet Lond. Engl. 395, 497–506 (2020).

4. Liao, M. et al. Single-cell landscape of bronchoalveolar immune cells in patients with COVID-19. Nat. Med. 26, 842–844 (2020).

5. Hu, B., Huang, S. & Yin, L. The cytokine storm and COVID-19. J. Med. Virol. 93, 250–256 (2021).

6. CDC. Types of COVID-19 Treatment. COVID-19 https://www.cdc.gov/covid/treatment/index.html (2024).

7. Focosi, D. et al. Monoclonal antibody therapies against SARS-CoV-2. Lancet Infect. Dis. 22, e311–e326 (2022).

8. Focosi, D. & Maggi, F. Neutralising antibody escape of SARS-CoV-2 spike protein: Risk assessment for antibody-based Covid-19 therapeutics and vaccines. Rev. Med. Virol. 31, e2231 (2021).

9. Errico, J. M., Adams, L. J. & Fremont, D. H. Antibody-mediated immunity to SARS-CoV-2 spike. Adv. Immunol. 154, 1–69 (2022).

10. Schoeman, D. & Fielding, B. C. Coronavirus envelope protein: current knowledge. Virol. J. 16, 69 (2019).

11. Xia, B. et al. SARS-CoV-2 envelope protein causes acute respiratory distress syndrome (ARDS)-like pathological damages and constitutes an antiviral target. Cell Res. 31, 847–860 (2021).

12. Zhou, S. et al. SARS-CoV-2 E protein: Pathogenesis and potential therapeutic development. Biomed. Pharmacother. Biomedecine Pharmacother. 159, 114242 (2023).

13. Troyano-Hernáez, P., Reinosa, R. & Holguín, Á. Evolution of SARS-CoV-2 Envelope, Membrane, Nucleocapsid, and Spike Structural Proteins from the Beginning of the Pandemic to September 2020: A Global and Regional Approach by Epidemiological Week. Viruses 13, 243 (2021).

14. Vann, K. R. et al. Binding of the SARS-CoV-2 envelope E protein to human BRD4 is essential for infection. Struct. Lond. Engl. 1993 30, 1224–1232.e5 (2022).

15. Caillet-Saguy, C. et al. Host PDZ-containing proteins targeted by SARS-CoV-2. FEBS J. 288, 5148–5162 (2021).

16. Ávila-Flores, A. et al. Identification of Host PDZ-Based Interactions with the SARS-CoV-2 E Protein in Human Monocytes. Int. J. Mol. Sci. 24, 12793 (2023).

17. Liu, L. et al. Coronavirus envelope protein activates TMED10-mediated unconventional secretion of inflammatory factors. Nat. Commun. 15, 8708 (2024).

18. Planès, R., Bert, J.-B., Tairi, S., BenMohamed, L. & Bahraoui, E. SARS-CoV-2 Envelope (E) Protein Binds and Activates TLR2 Pathway: A Novel Molecular Target for COVID-19 Interventions. Viruses 14, 999 (2022).

19. Teoh, K.-T. et al. The SARS coronavirus E protein interacts with PALS1 and alters tight junction formation and epithelial morphogenesis. Mol. Biol. Cell 21, 3838–3852 (2010).

20. Shepley-McTaggart, A. et al. SARS-CoV-2 Envelope (E) Protein Interacts with PDZ-Domain-2 of Host Tight Junction Protein ZO1. BioRxiv Prepr. Serv. Biol. 2020.12.22.422708 (2020) doi:10.1101/2020.12.22.422708.

21. Jimenez-Guardeño, J. M. et al. The PDZ-binding motif of severe acute respiratory syndrome coronavirus envelope protein is a determinant of viral pathogenesis. PLoS Pathog. 10, e1004320 (2014).

22. Alvarez, F. et al. The SARS-CoV-2 envelope PDZ Binding Motif acts as a virulence factor disrupting host’s epithelial cell-cell junctions. 2024.12.10.627528 Preprint at 10.1101/2024.12.10.627528 (2024).

23. Honrubia, J. M. et al. SARS-CoV-2-Mediated Lung Edema and Replication Are Diminished by Cystic Fibrosis Transmembrane Conductance Regulator Modulators. mBio 14, e0313622 (2023).

24. Zhu, Y. et al. Interactions of Severe Acute Respiratory Syndrome Coronavirus 2 Protein E With Cell Junctions and Polarity PSD-95/Dlg/ZO-1-Containing Proteins. Front. Microbiol. 13, 829094 (2022).

25. Javorsky, A., Humbert, P. O. & Kvansakul, M. Structural basis of coronavirus E protein interactions with human PALS1 PDZ domain. *Commun*. Biol. 4, 724 (2021).

26. Chai, J. et al. Structural basis for SARS-CoV-2 envelope protein recognition of human cell junction protein PALS1. Nat. Commun. 12, 3433 (2021).

27. Deinhardt-Emmer, S. et al. SARS-CoV-2 causes severe epithelial inflammation and barrier dysfunction. J. Virol. 95, e00110–21, JVI.00110-21 (2021).

28. Robinot, R. et al. SARS-CoV-2 infection induces the dedifferentiation of multiciliated cells and impairs mucociliary clearance. Nat. Commun. 12, 4354 (2021).

29. Lamers, M. M. & Haagmans, B. L. SARS-CoV-2 pathogenesis. Nat. Rev. Microbiol. 20, 270–284 (2022).

30. Xu, J.-B. et al. SARS-CoV-2 envelope protein impairs airway epithelial barrier function and exacerbates airway inflammation via increased intracellular Cl- concentration. Signal Transduct. Target. Ther. 9, 74 (2024).

31. Bosc, N. et al. Fr-PPIChem: An Academic Compound Library Dedicated to Protein-Protein Interactions. ACS Chem. Biol. 15, 1566–1574 (2020).

32. Fanning, A. S., Lye, M. F., Anderson, J. M. & Lavie, A. Domain swapping within PDZ2 is responsible for dimerization of ZO proteins. J. Biol. Chem. 282, 37710–37716 (2007).

33. Chen, J., Pan, L., Wei, Z., Zhao, Y. & Zhang, M. Domain-swapped dimerization of ZO-1 PDZ2 generates specific and regulatory connexin43-binding sites. EMBO J. 27, 2113–2123 (2008).

34. Zhang, J. H., Chung, T. D. & Oldenburg, K. R. A Simple Statistical Parameter for Use in Evaluation and Validation of High Throughput Screening Assays. J. Biomol. Screen. 4, 67– 73 (1999).

35. Lang, P. T. et al. DOCK 6: Combining techniques to model RNA–small molecule complexes. RNA 15, 1219–1230 (2009).

36. Gupta, A. et al. Extrapulmonary manifestations of COVID-19. Nat. Med. 26, 1017– 1032 (2020).

37. Chen, Y., Yang, W., Chen, F. & Cui, L. COVID-19 and cognitive impairment: neuroinvasive and bloodlJbrain barrier dysfunction. J. Neuroinflammation 19, 222 (2022).

38. Schnittler, H. J. Structural and functional aspects of intercellular junctions in vascular endothelium. Basic Res. Cardiol. 93 **Suppl 3**, 30–39 (1998).

39. Chen, C.-H. et al. The connexin 43/ZO-1 complex regulates cerebral endothelial F-actin architecture and migration. Am. J. Physiol. Cell Physiol. 309, C600–607 (2015).

40. Weber, P. et al. High-Throughput Crystallization Pipeline at the Crystallography Core Facility of the Institut Pasteur. Mol. Basel Switz. 24, 4451 (2019).

41. Kabsch, W. XDS. Acta Crystallogr. D Biol. Crystallogr. 66, 125–132 (2010).

42. McCoy, A. J. Solving structures of protein complexes by molecular replacement with Phaser. Acta Crystallogr. D Biol. Crystallogr. 63, 32–41 (2007).

43. Emsley, P., Lohkamp, B., Scott, W. G. & Cowtan, K. Features and development of Coot. Acta Crystallogr. D Biol. Crystallogr. 66, 486–501 (2010).

44. Adams, P. D. et al. PHENIX: a comprehensive Python-based system for macromolecular structure solution. Acta Crystallogr. D Biol. Crystallogr. 66, 213–221 (2010).

45. Gally, J.-M., Bourg, S., Do, Q.-T., Aci-Sèche, S. & Bonnet, P. VSPrep: A General KNIME Workflow for the Preparation of Molecules for Virtual Screening. Mol. Inform. 36, (2017).

46. Gally, J.-M. et al. VSPrep: A KNIME Workflow for the Preparation of Molecular Databases for Virtual Screening. Curr. Med. Chem. 27, 6480–6494 (2020).

47. Hamon, V. et al. 2P2IHUNTER: a tool for filtering orthosteric protein–protein interaction modulators via a dedicated support vector machine. J. R. Soc. Interface 11, 20130860 (2014).

48. Baell, J. B. & Holloway, G. A. New substructure filters for removal of pan assay interference compounds (PAINS) from screening libraries and for their exclusion in bioassays. J. Med. Chem. 53, 2719–2740 (2010).

49. Glem, R. C. et al. Circular fingerprints: flexible molecular descriptors with applications from physical chemistry to ADME. IDrugs Investig. Drugs J. 9, 199–204 (2006).

50. Butina, D. Unsupervised Data Base Clustering Based on Daylight’s Fingerprint and Tanimoto Similarity: A Fast and Automated Way To Cluster Small and Large Data Sets. J. Chem. Inf. Comput. Sci. 39, 747–750 (1999).

51. Pettersen, E. F. et al. UCSF Chimera--a visualization system for exploratory research and analysis. J. Comput. Chem. 25, 1605–1612 (2004).

52. O’Boyle, N. M. et al. Open Babel: An open chemical toolbox. J. Cheminformatics 3, 33 (2011).

53. de Melo, G. D. et al. Neuroinvasion and anosmia are independent phenomena upon infection with SARS-CoV-2 and its variants. Nat. Commun. 14, 4485 (2023).

